# A stress-response-related inter-compartmental signalling pathway regulates embryonic cuticle integrity in Arabidopsis

**DOI:** 10.1101/477109

**Authors:** Audrey Creff, Lysiane Brocard, Jérôme Joubès, Ludivine Taconnat, Nicolas M. Doll, Stephanie Pascal, Roberta Galletti, Anne-Charlotte Marsollier, Steven Moussu, Thomas Widiez, Frédéric Domergue, Gwyneth Ingram

## Abstract

The embryonic cuticle is necessary for normal seed development and seedling establishment in Arabidopsis. Although mutants with defective embryonic cuticles have been identified, neither the deposition of cuticle material, nor its regulation, has been described during embryogenesis. Here we use electron microscopy, lipid staining and permeability assays to show that cuticle deposition initiates *de novo* in patches on globular embryos. By combining these techniques with genetics and gene expression analysis, we show that successful patch coalescence to form a continuous cuticle requires a signalling involving the endosperm-specific subtilisin protease ALE1 and the receptor kinases GSO1 and GSO2, which are expressed in the developing embryonic epidermis. Transcriptome analysis shows that this pathway regulates stress-related gene expression in seeds. Consistent with these findings we show genetically, and through activity analysis, that the stress-associated MPK6 protein acts downstream of GSO1 and GSO2 in the developing embryo. We propose that a stress-related signalling pathway has been hijacked in some angiosperm seeds through the recruitment of endosperm-specific components. Our work reveals the presence of an inter-compartmental dialogue between the endosperm and embryo that ensures the formation of an intact and functional cuticle around the developing embryo through an “auto-immune” type interaction.

## INTRODUCTION

The Arabidopsis seed is a complex structure composed of three genetically distinct compartments, the maternally-derived seed coat, the embryo, and the endosperm. After fertilization the expansion of the endosperm drives the growth of the seed. However, during later developmental stages the endosperm breaks down, leaving space for the growing embryo. By the end of seed development, only a single endosperm cell layer envelops the embryonic tissues (reviewed in [1]).

The endosperm is an angiosperm innovation, thought to have arisen through the sexualisation of the central cell of the female gametophyte [2]. The ancestors of angiosperms probably had seeds more similar to those of gymnosperms, in which tissues of the female gametophyte proliferate independently of egg cell fertilization to produce a nutrient rich storage tissue. However, the endosperm plays not only a nutritional role, but also a role in regulating embryo development. For example, the peptide CLAVATA3/EMBRYO SURROUNDING REGION-RELATED8 (CLE8) may act non-cell autonomously to regulate early Arabidopsis embryogenesis [3]. Recently, maternally-expressed peptides present in the central cell pre-fertilization, and subsequently in the early EMBRYO SURROUNDING REGION (ESR), were shown to regulate Arabidopsis suspensor development. Genetic analysis suggests that this regulation could be mediated by a pathway involving the Receptor-Like Cytoplasmic Kinase SHORT SUSPENSOR [4,5], although the receptor involved remains unidentified.

In previous works we showed genetically that the ESR-specific subtilisin protease Abnormal LEaf-shape1 (ALE1) acts in the same genetic pathway as two embryonically-expressed receptor kinases, GASSHO1 [(GSO1) also known as SCHENGEN3 [6]] and GASSHO2 (GSO2), to control the formation of the embryonic cuticle in developing seeds [7-10]. Our results indicate that a seed specific inter-tissue signalling event is necessary for the formation of a functional embryonic cuticle [7]. The results of genetic studies have led us to speculate that the role of this pathway is to ensure the robust elimination of apoplastic continuity between the developing embryo and the surrounding endosperm thus gating molecular movement between the two compartments [11,12].

The cuticle is the outermost layer of the aerial parts of the plant. It is a highly complex structure mainly composed of a lipid polymer (cutin) and waxes, either associated with the polymer (intracuticular waxes) or deposited on the top of it (epicuticular waxes) (recently reviewed in [13,14]). Cutin and waxes are composed of complex mixtures of hydroxylated and very long-chain fatty acid derivatives, respectively. Cuticle structure and composition are highly regulated not only at the tissue level, but also in response to environmental stimuli such as drought, radiation and pollution [13,14]. In addition, several reports have highlighted the important role played by the cuticle in biotic interactions, and particularly in protecting plants from attack by bacterial pathogens (reviewed in [13,15,16]).

In Arabidopsis although little, if any, evidence exists for the presence of cutin-like substances in the wall between the mature egg cell and the central cell, by the end of embryogenesis the hypocotyl and cotyledons of the embryo are covered with a continuous cuticle which renders the germinating seedling impermeable to hydrophilic dyes, and resistant to water loss [17]. Cuticle biogenesis is considered to be a unique property of epidermal cells [18]. During plant development, epidermal cells are generated by anticlinal divisions of pre-existing epidermal cells so that each cell inherits an intact external cuticularised cell wall. In this respect the embryonic cuticle is atypical as it is deposited *de novo* at the interface between the developing embryo and endosperm. Although mutants with defective embryonic cuticles have been described [7-10,17], only very fragmentary evidence about when the embryonic cuticle appears is present in the literature. Furthermore, the structure of the embryonic cuticle, its composition, the mechanisms via which it is deposited and its function during seed development remain unexplored. In this study we aimed to elucidate how the embryonic cuticle is formed, and to investigate how the ALE1 GSO1 GSO2 signalling pathway impacts its biosynthesis and deposition.

## RESULTS

### Expression of genes involved in cuticle deposition initiates during early embryogenesis

An inspection of available *in silico* data [19-21] showed that many genes encoding enzymes thought to be involved in cutin biosynthesis are expressed during early embryogenesis (Supplementary Figure 1). *In situ* hybridisations confirmed that genes known to affect cuticle production (*LACS2* [22,23], *FIDDLEHEAD/KCS10* [24-27], *LACERATA* [28] and *BODYGUARD* [29]) or export (*LTPG1* [30] and *ABCG11* [31]) have a clear epidermis-specific expression from the mid globular stage onwards (Figure 1, Supplementary Figure 2, Supplementary Figure 3).

**Figure 1.**
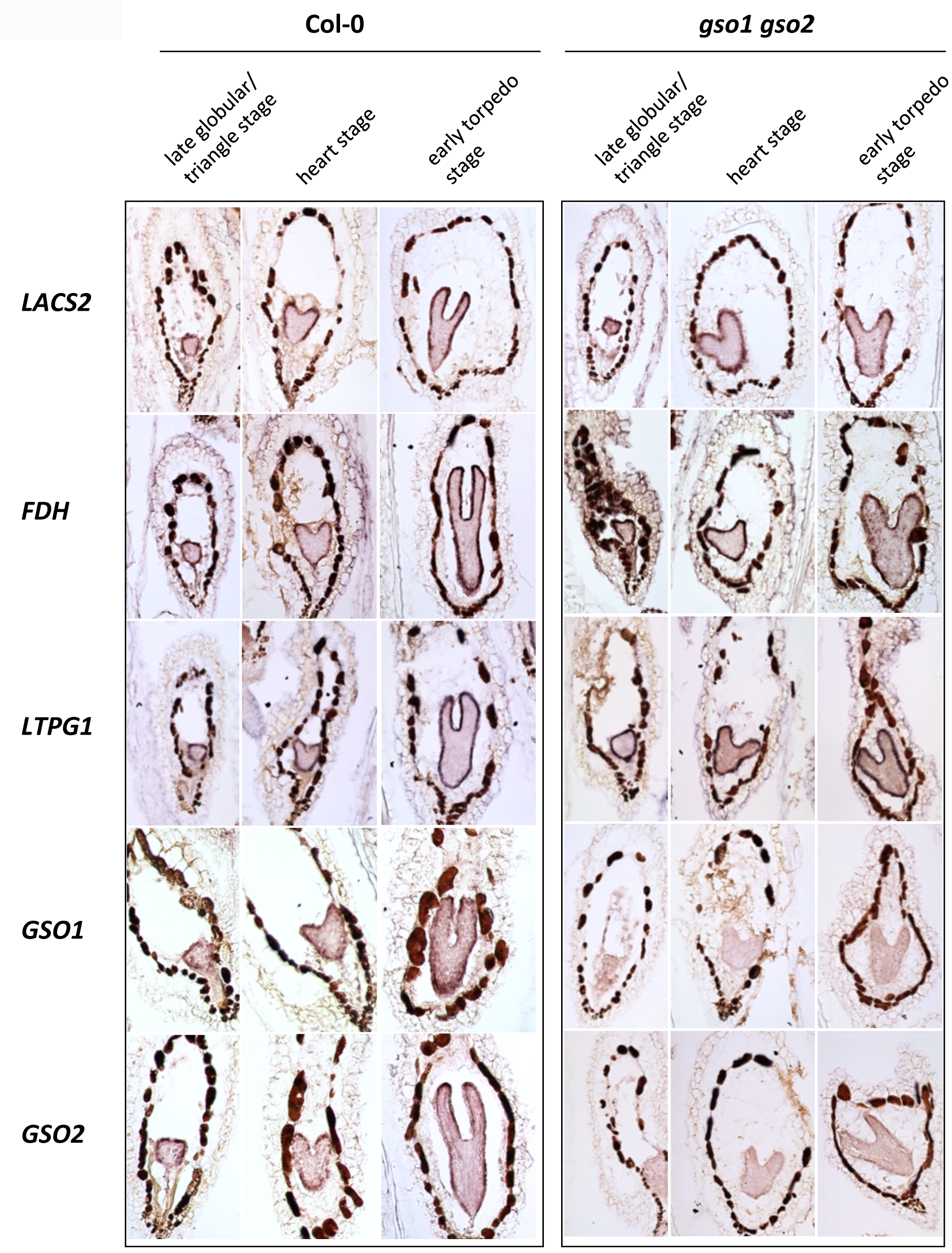
Genes involved in cuticle biosynthesis are co-expressed with *GSO1* and *GSO2* during embryogenesis, but their expression is not dependent upon GSO1 and GSO2. Analysis of the expression of genes involved in cuticle biosynthesis in wild-type (Col-0) and *gso1-1 gso2-1* seeds containing late globular/triangle, heart and early torpedo stage embryos (left to right).

In agreement with published and *in silico* data [9] (Supplementary Figure 1) *GSO1* and *GSO2* were expressed in the embryo from early developmental stages (Figure 1, Supplementary Figure 3). In addition, their expression was mainly restricted to the embryonic epidermis. *GSO1* expression in the embryonic epidermis was further confirmed using plants expressing a functional genomic GSO1-mVENUS fusion under the control of the *GSO1* promoter (*pGSO1*:*GSO1-mVENUS*) [6] (Figure 2). This construction fully complemented the cuticle permeability phenotype of *gso1-1 gso2-1* double mutant seedlings, and strongly reduced the misshapen-seed phenotype of *gso1-1 gso2-1* mutant seeds when introduced into the *gso1-1 gso2-1* mutant background (Figure 2).

**Figure 2.**
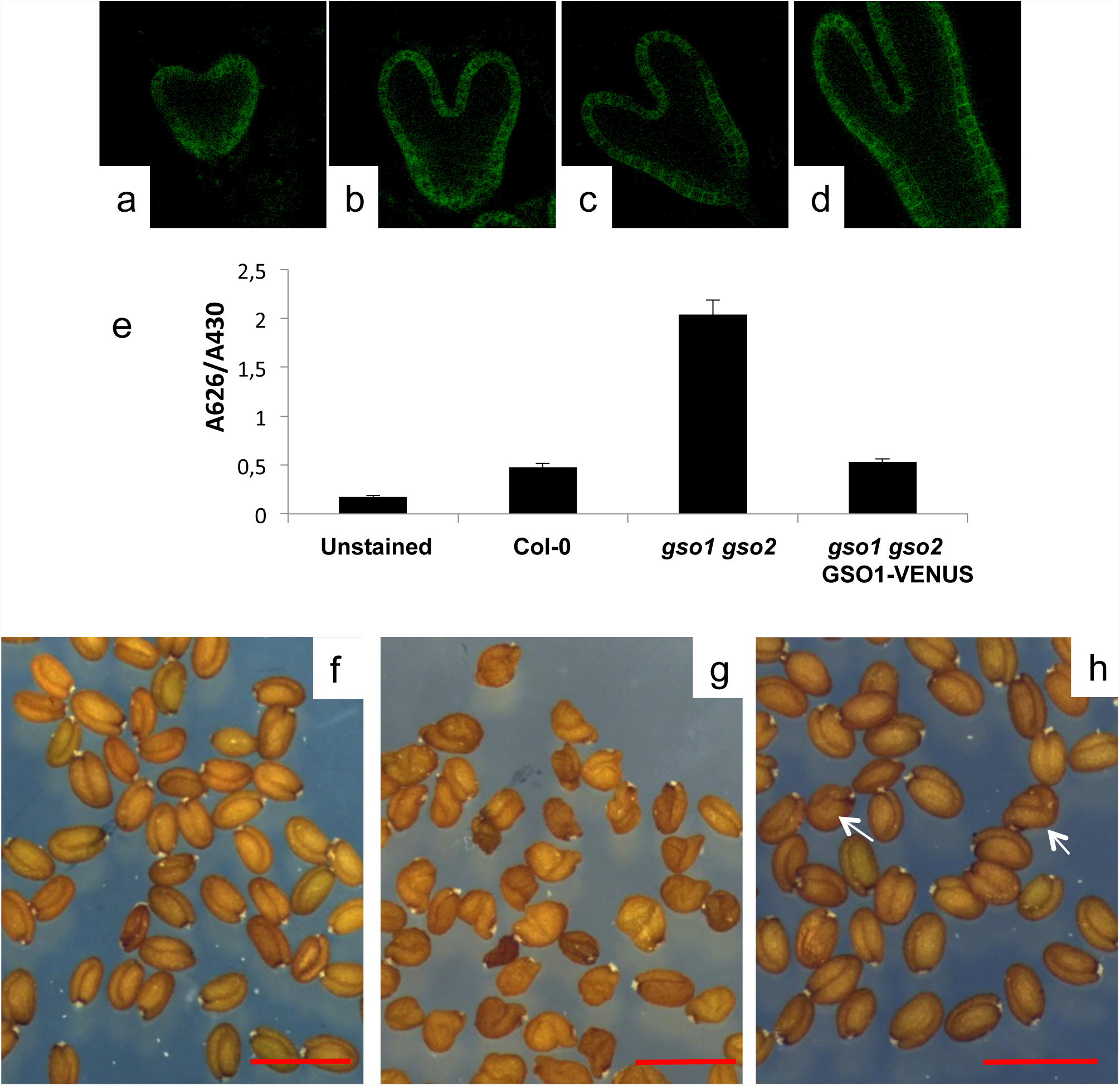
Localisation and functionality of the *pGSO1:GSO1-mVENUS transgene* in developing embryos. Confocal images of GSO1-mVENUS at early heart, mid heart, late heart and early torpedo stages of development (a-d). (e) The *pGSO1:GSO1-mVENUS* transgene complements seedling cu6cle defects in *gso1-1 gso2-1* double mutants. Quantification of seedling toluidine blue permeability was carried out as described in Moussu et al., 2017. Error bars represent standard errors from three biological replicates. (f-h) The *pGSO1:GSO1-mVENUS* transgene complements seed shape defects in *gso1-1 gso2-1* double mutants. Seed populations from wild-type (f), *gso1-1 gso2-1* double mutants (g) and *gso1-1 gso2-1* double mutants carrying the The *pGSO1:GSO1-mVENUS* transgene (h). Occasional misshapen shaped seeds are observed in the complemented line (white arrows), compared with 100% misshapen seeds in the un-complemented double mutant. Scale bar = 1mm.

Since previous results showed that epidermal identity is not affected in *gso1-1 gso2-1* mutants [12], the expression of cuticle biosynthesis genes was analysed by *in situ* hybridization in *gso1-1 gso2-1* double mutant seeds (which show a stronger cuticle phenotype than *ale1-4* mutants [7]). As shown in Figure 1 (and Supplementary Figure 2, Supplementary Figure 3), no reduction in the expression of any of the cuticle biogenesis genes analysed was detected in the embryonic epidermis of this background, whereas reduced expression of both *GSO1* and *GSO2* was clearly visible. For these results, we concluded that although many genes involved in cuticle biosynthesis are co-expressed with GSO1 and GSO2 in the embryonic epidermis, their expression is not dependent upon GSO1 and GSO2.

### Loss of GSO1/GSO2 and ALE1 function affects cuticle integrity

The cutin content of seedling cotyledons was assessed by measuring the quantities of the main cutin monomers released after cutin isolation followed by depolymerisation (mainly C16 and C18 ωOH (omega-hydroxy acid) and DCA (α,ω-dicarboxylic acid)). As clearly illustrated by the quantification of 18:2-DCA, the major component of Arabidopsis cutin, a slight loss in cutin load was detected in *gso1-1 gso2-1*, but not in *ale1-4* cotyledons compared to wild-type. In contrast a very clear reduction in cutin load was observed in control plants lacking the acyltransferases GPAT4 and GPAT8 required for cutin biosynthesis, as has previously been reported in rosette leaves [32] (Figure 3a). We therefore investigated the cuticle permeability of etiolated cotyledons by submerging them in the hydrophilic dye toluidine blue, which can only penetrate internal tissues through defects in the cuticle [17]. Surprisingly, we found that the cotyledons of etiolated *gpat4 gpat8* seedlings showed a rather similar toluidine blue permeability to *ale1-4* seedlings and a considerably reduced permeability compared to *gso1-1 gso1-2* double mutants, suggesting that the *gpat4 gpat8* cuticle, although quantitatively strongly deficient in cutin monomers, remains partially functional (Figure 3b). Taken together with gene expression analysis, these results suggest that the ALE1, GSO1 and GSO2-mediated signalling pathway might impact cuticle organisation or integrity rather than the quantity of cuticle components produced by epidermal cells.

**Figure 3.**
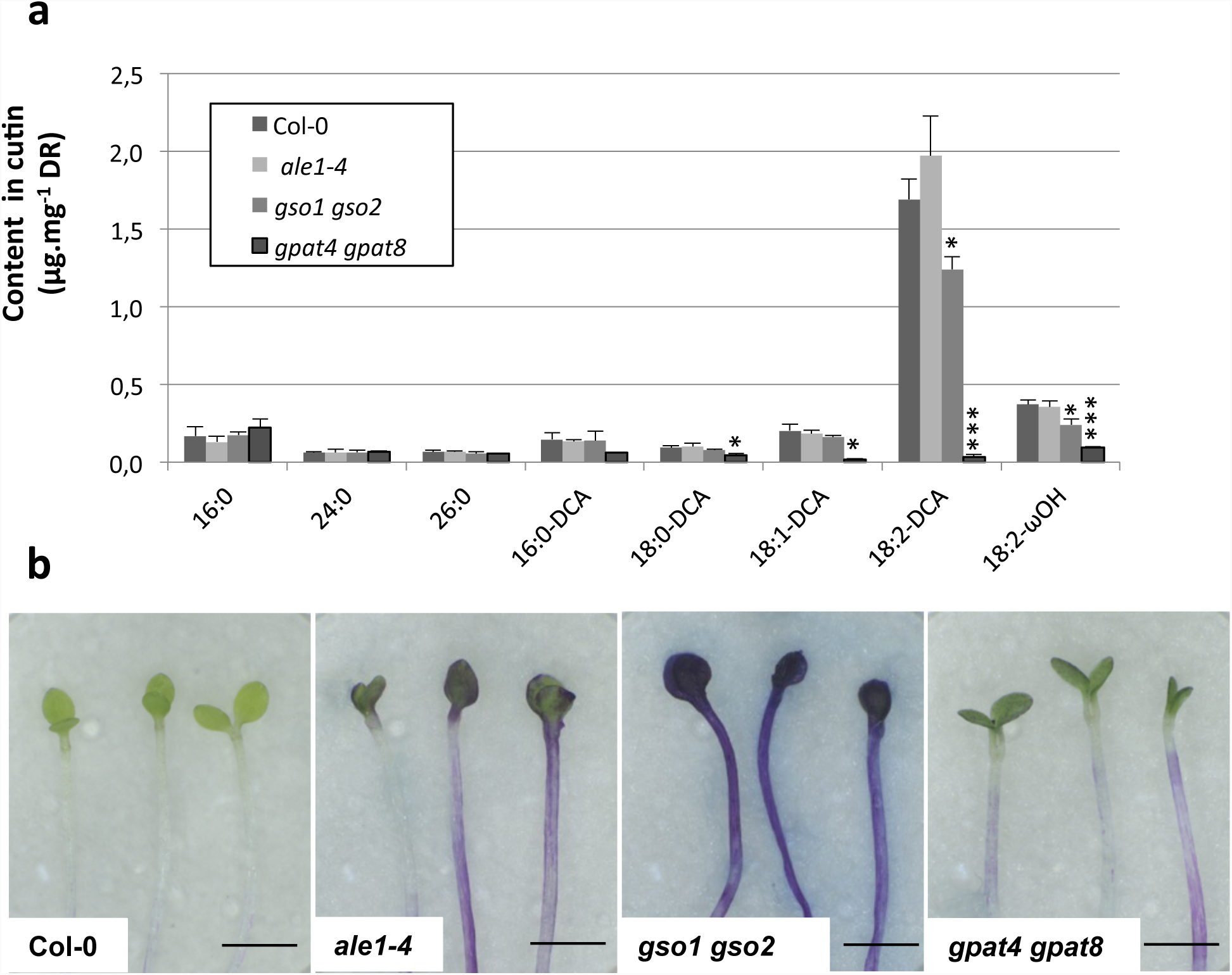
Cuticle permeability defects in *ale1-4* and *gso1-1 gso2-1* seedlings do not correlate with changes in cutin load. (a) Cotyledons grown *in vitro* for 5 days under continuous light were collected, delipidated and their cutin content and composition was analyzed as described in the Material and Method section. ωOH and DCA stand respectively for omega-hydroxy acid and α,ω-dicarboxylic acid. Mean values are shown in μg/mg of delipidated dry residue (DR) ± SD of three replicates. Statistical differences were determined according to a Student’s *t* test : *** denotes p<0.0001, ** denotes p<0.001 and * denotes p<0.01. (b) Cuticle permeability to toluidine blue in etiolated seedlings from the genotypes tested in (a). Scale bar = 2mm.

The process of embryonic cuticle deposition was investigated in more detail in wild-type (Col-0) seeds (Figure 4a-d). At the two-cell stage the embryo was surrounded by a thick cell wall but no electron dense material was detected at the embryo surface. At the mid-late globular stage, a cutin-like electron-dense material was detected in patches (Figure 4b and Supplementary Figure 5a,b). From heart stage onwards, an apparently continuous layer of electron-dense cutin-like material was detected at the surface of the outer epidermal cell wall. Embryonic cuticle production therefore involves the *de novo* deposition and subsequent coalescence of “patches” of cuticular material at the surface of epidermal cells. Toluidine blue assays with wild-type embryos extruded at different developmental stages indicated that permeability started to reduce noticeably at the early torpedo stage (slightly after apparent gap closure), and that the embryo continued to become more and more impermeable during embryo development (Supplementary Figure 4), suggesting that the coalescence of gaps in the embryonic cuticle correlates well with a reduction in embryonic permeability.

**Figure 4.**
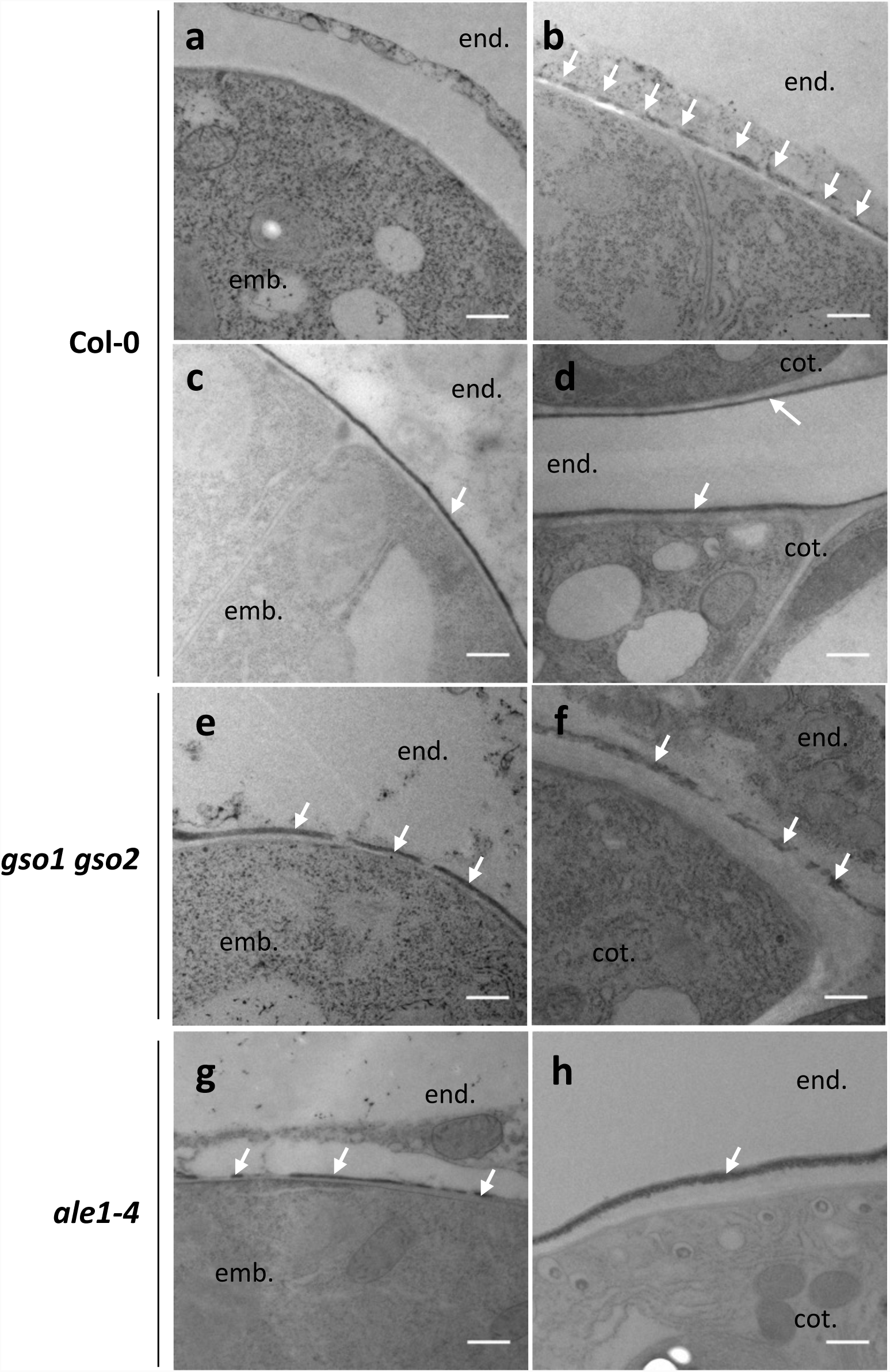
Embryonic cuticle biogenesis involves a process of patch coalescence that is defective in *ale1-4* and *gso1-1 gso2-1* mutants. Analysis of embryonic cuticle deposi6on in wild-type (a-d), *gso1-1 gso2-1 (*e-f) and *ale1-4 (*g,h) embryos at 2 cell (a), mid globular (b), mid heart (c,e,g) and walking s6ck (d,f,h) stages of embryogenesis. White arrows show the external face of the embryonic cuticle. (Embryo (emb.), endospem (end.) and cotyledon (cot.) (for late stages), are labelled. Scale bar = 500nm.

In *gso1-1 gso2-1* mutants the cuticle still showed discontinuities at the heart and walking stick stage (Figure 4e-f, Supplementary Figure 5c-f). In this background the cuticle also appeared thicker, but less condensed than that of wild-type embryos. The outer epidermal cell wall was also abnormally thick at later stages (compare embryonic cell wall thickness in Figure 4d with that in 4f). Similar discontinuities were observed, although at a lower frequency, in the *ale1-4* background at the heart stage as described previously [10], but were less frequent at later stages, consistent with the less severe cuticle permeability phenotype observed in the seedlings of this background (Figure 4g-h). These results are consistent with our hypothesis that the ALE1 GSO1 GSO2 pathway is necessary for generating a continuous cuticle layer and further suggest that it controls “gap closure” during embryonic cuticle maturation.

### GSO1 GSO2 and ALE1 regulate overlapping gene sets and promote the expression of defence related genes during seed development

Transcriptional analysis of intact siliques from *gso1-1 gso2-1* and *ale1-4* mutants and wild-type plants was carried out at globular and heart stages. The results are provided in Figure 5a and b, Supplementary Table 1, and Supplementary Figures 6 and 7. The number of differentially down-regulated genes in the mutant backgrounds compared to wild-type was higher than the number of up-regulated genes (Supplementary Table 1). A moderate overlap between genes showing higher expression in *ale1-4* and *gso1-1 gso2-1* mutants than wild-type controls was observed (Supplementary Figure 6, Supplementary Table 1). In contrast more than three quarters of the genes showing reduced expression at both developmental stages in the *gso1-1 gso2-1* background also showed reduced expression at both developmental stages in *ale1-4* mutants (Figure 5a and Supplementary Table 1), corroborating previously published genetic evidence that ALE1, GSO1 and GSO2 act in the same genetic pathway [7]. Because *ALE1* appears to be expressed exclusively in the ESR region of the endosperm [7,8,10], genes mis-regulated in both mutant backgrounds likely comprise *bona fide* targets (direct and indirect) of the ALE1 GSO1 GSO2 pathway, despite the fact that the expression of *GSO1* and *GSO2* is not restricted to the seed [6,9,20].

**Figure 5.**
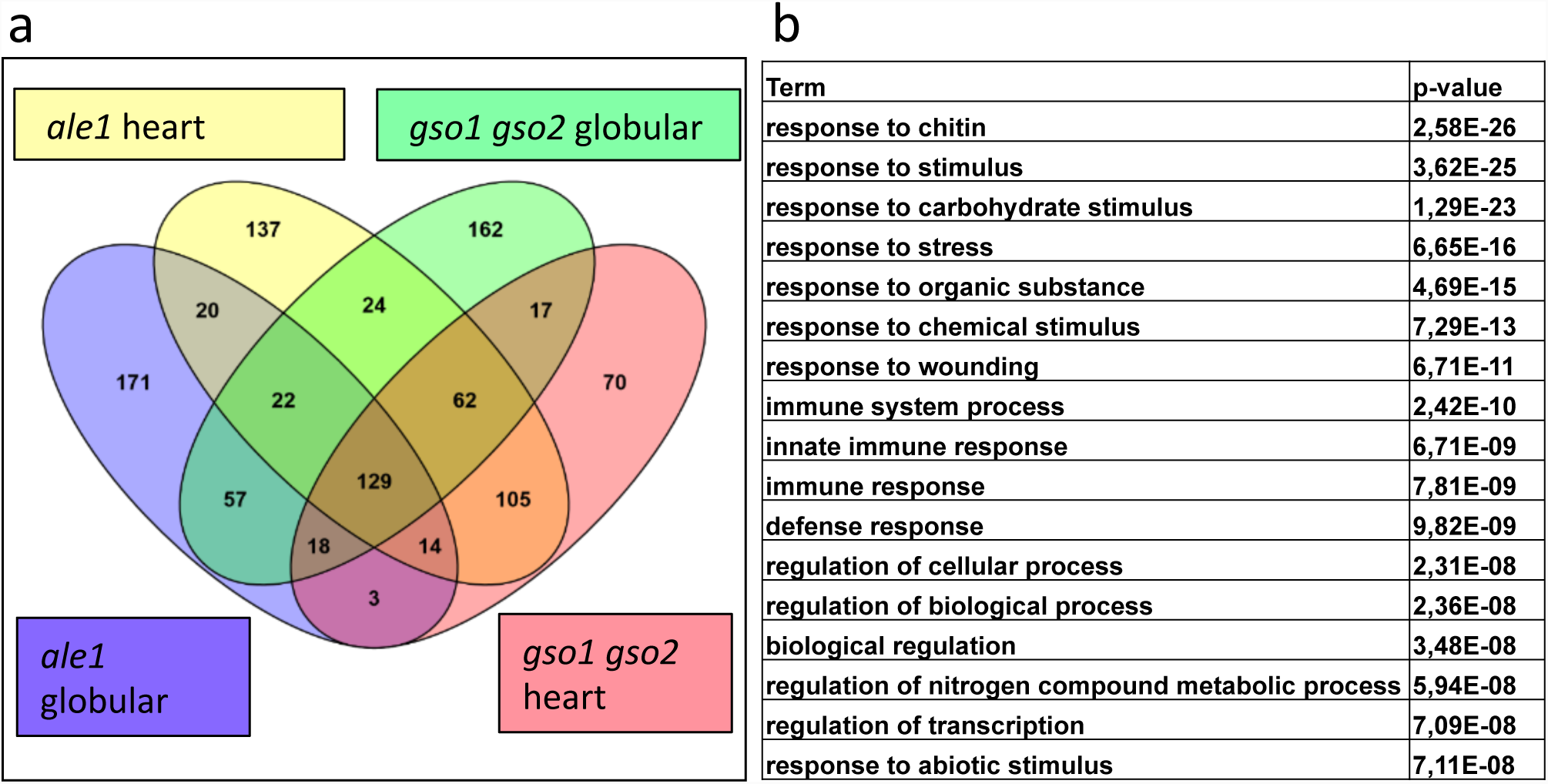
ALE1, GSO1 and GSO2 positively regulate the expression of stress-related genes in seeds. (a) Summary of overlaps between gene sets showing reduced expression in *ale1-4* and *gso1-1 gso2-1* mutants at the globular and heart stages of development. (b) GO term analysis of genes down-regulated in both *ale1-4* and *gso1-1 gso2-2* mutants at the heart stage.

Genes up-regulated in both mutant backgrounds showed a moderate over-representation in GO terms associated with responses to abiotic stress (Supplementary Figure 6). In contrast, genes down-regulated in both backgrounds, particularly at the heart stage, showed a very striking overrepresentation for GO terms linked to abiotic and biotic stress responses (Figure 5b, Supplementary Figure 7). Mis-regulation of 19 of these genes was validated using additional independent biological samples by qRT-PCR (Supplementary Figure 8). The expression levels of these genes in seeds were generally low, and attempts to carry out *in situ* hybridization were inconclusive. However for one target, *SWI3A* [33], expression in the developing embryo predicted from *in silico* data was confirmed, and shown to be convincingly reduced in embryos of the *gso1-1 gso2-1* double mutant (Supplementary Figure 9). Thus, consistent with the embryonic expression of *GSO1* and *GSO2*, some of the transcriptional regulation downstream of ALE1 GSO1 GSO2 signalling occurs in the embryo. Expression of *ALE1* was not reduced in *gso1-1 gso2-1* mutants (Supplementary Table 2 and Supplementary Figure 8), suggesting that ALE1 is not a downstream target of GSO1 GSO2-mediated signalling, and could therefore act upstream of GSO1 and GSO2 in mediating embryonic responses necessary for the establishment of an intact embryonic cuticle.

### MPK6 acts in the ALE1 GSO1 GSO2 signalling pathway

The GSO1 and GSO2 receptor kinases belong to family XI of the Leucine-Rich Repeat (LRR)-RLKs [34,35], and are closely related to the “danger” peptide receptors PEPR1 and PEPR2 [36,37], which are involved in the amplification of defence responses triggered by pathogen-associated molecular pattern (PAMP) perception [38]. A previous study [39], reported aberrantly shaped seeds, resembling those of *ale1-4* mutants, in Arabidopsis *mpk6* mutants lacking the MITOGEN ACTIVATED PROTEIN KINASE6 (MPK6) protein, which acts downstream of PEPR signalling. In addition a proportion of *mpk6* mutant seeds were reported to rupture [39]. We confirmed these phenotypes in the *mpk6-2* mutant background (Supplementary Figure 10). A recent article has suggested that some seed defects in *mpk6* mutants may depend upon the genotype of the maternal tissues in the seed [40]. Reciprocal crosses were therefore performed, and these confirmed that seed twisting phenotype is dependent upon the genotype of the zygotic compartment and not the maternal compartment (Supplementary Figure 11). We found that a proportion of *mpk6-2* seedlings showed abnormal permeability to the hydrophilic dye toluidine blue, consistent with the presence of cuticle defects (Figure 6). Nile red staining [32] of the cotyledons of etiolated seedlings was used to confirm *mpk6* cuticle defects (Supplementary Figure 12 a,b). Using this technique, wild-type cotyledons were found to be covered with a continuous lipid cuticle layer. As previously reported, and consistent with our cutin analysis, *gpat4 gpat8* mutants showed drastically reduced cuticle staining. In contrast *gso1-1 gso2-1* mutants showed a patchy cuticle, similar to that seen using transmission electron microscopy on the embryo surface. Both *ale1-4* and *mpk6-2* mutants showed a less well-defined cuticle than wild-type, which although apparently continuous, showed uneven cutin deposition (Supplementary Figure 12 a,b).

**Figure 6.**
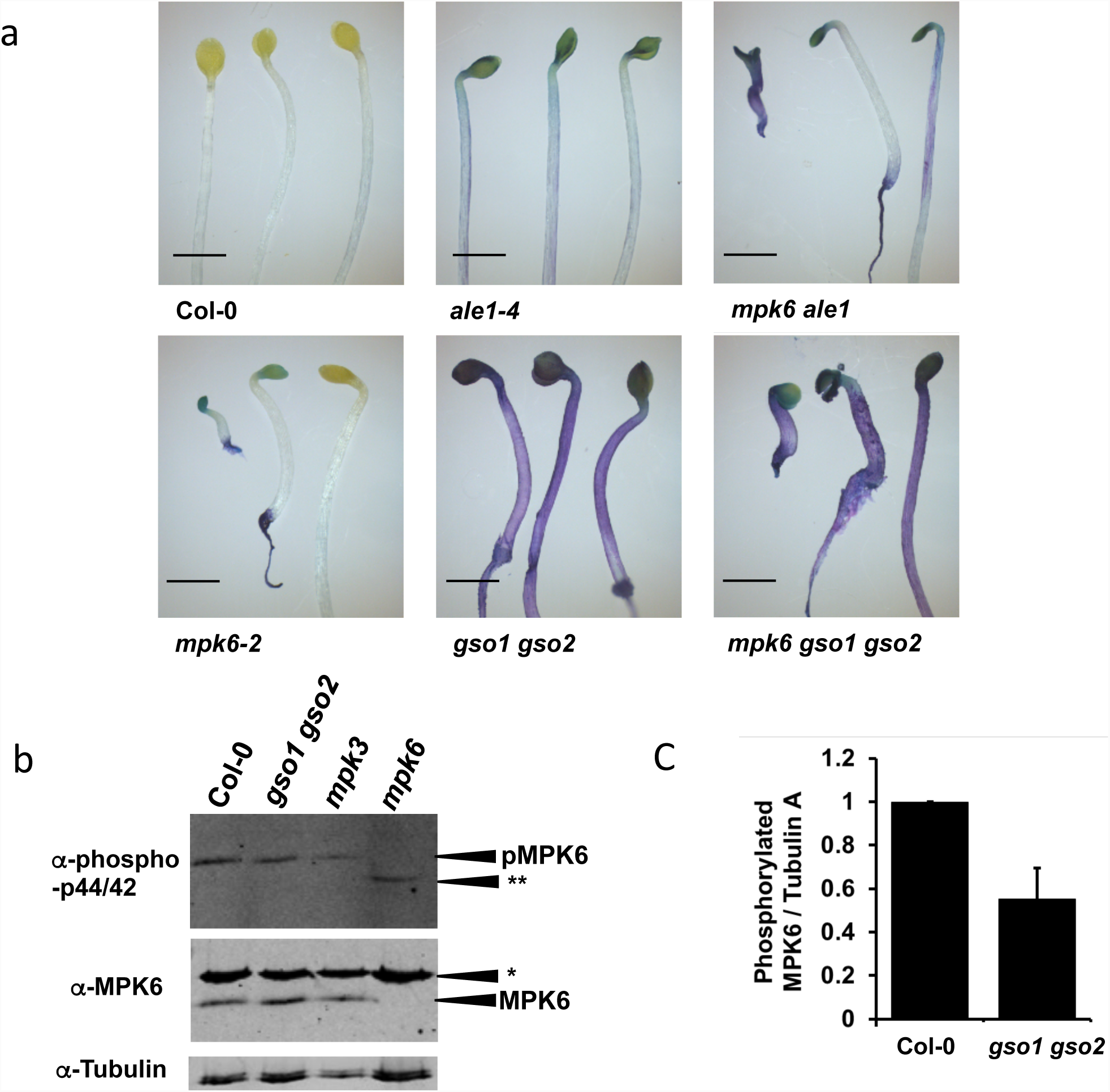
MPK6 acts downstream of ALE1, GSO1 and GSO2 mediated signalling. (a) Seedling cu6cle permeability phenotypes of *mpk6-2* and *ale1-4* and *gso1-1 gso2-1* mutants and in double and triple mutant combinations. Scale bar = 2mm (b) Analysis of proteins extracted from developing seeds at the globular-early torpedo stage. The mutants *mpk3-1* and *mpk6-2* were included to confirm band identification. No phosphorylation of MPKs other than MPK6 is observed in Col-0, *gso1-1 gso2-1* or *mpk3-1* seeds, but an additional band (**) is systematically observed in the *mpk6-2* mutant background. * Indicates a non specific band detected by the anti-MPK6 antibody. This experiment was repeated 7 on independent biological samples, with similar results. (c) Degree of phosphoryla6on of MPK6 in Col-0 and *gso1-1 gso2-1* mutant seeds. Error bars represent SD of 3 biological replicates (see Supplementary Figure 16 for linearity testing).

Triple *mpk6-2 gso1-1 gso2-1* and double *ale1-4 mpk6-2* mutants were generated to investigate further the genetic interactions of *ALE1, GSO1* and *GSO2* with *MPK6.* Fertility in *ale1-4 mpk6-2* double mutants was similar to that in *mpk6-2* mutants, while triple *mpk6-2 gso1-1 gso2-1* mutant plants were viable but produced very few seeds. In terms of seed shape and cotyledon cuticle permeability, triple *mpk6-2 gso1-1 gso2-1* mutants had phenotypes identical to those observed in *gso1-1 gso2-1* double mutants (Figure 6, Supplementary Figure 10). Since all *gso1-1 gso2-1* mutant seeds are twisted, non-additivity cannot be concluded from this phenotype. However, recent work has shown that additivity of toluidine blue staining phenotypes can be detected in mutant combinations with *gso1-1 gso2-1 [41].* The frequency of “twisted” seeds (including ruptured seeds), and toluidine blue stained seedling cotyledons was non-additive in *ale1-4 mpk6-2* double mutant plants, consistent with *ALE1, GSO1, GSO2* and *MPK6* acting in the same genetic pathway to control seedling cotyledon permeability (Figure 6 and Supplementary Figure 10).

MPK6 is involved in a plethora of reproductive and non-reproductive developmental processes and shows functional redundancy with other MPK proteins [39,42-52] meaning that global transcriptome analysis in the *mpk6-2* background would likely be uninformative for this study. We therefore directly tested a subset of genes mis-regulated in *gso1-1 gso2-1* and *ale1-4* mutants for misregulation in *mpk6-2* mutants at three stages of embryo development. Five out of eight genes tested showed reduced expression in *mpk6-2* either at all three stages (*SWI3A, WRKY70* and *NIMIN1*), or in two out of three developmental stages tested (*SIB1* and *NIMIN2*) (Supplementary Figure 13). Unsurprisingly given the relatively weak cuticle phenotype of *mpk6* mutants compared with *gso1 gso2* mutants, some genes showing strong down-regulation in the *gso1-1 gso2-1* mutants (*WRKY33, WRKY46 and WRKY53)* did not show any significant reduction in expression in the *mpk6-2* mutant background (Supplementary Figure 13) indicating that their transcriptional regulation downstream of GSO1 and GSO2-mediated signalling could be dependent on signalling components acting redundantly with MPK6. The expression of *ALE1, GSO1* and *GSO2* was not altered in *mpk6-2* mutants (Supplementary Figure 13), indicating that MPK6 most probably acts downstream of GSO1 and GSO2-mediated signalling.

To further confirm this hypothesis, we analysed MPK phosphorylation in developing seeds from Col-0 and *gso1-1 gso2-1* double mutants. In seedlings, phosphorylation of MPK6 (and additional MPKs) can only be detected after elicitation (for example with the flg22 peptide). The response to flg22 is not attenuated in *gso1/gso2* mutant seedlings (Supplementary Figure 14 and 15). In contrast, MPK6 phosphorylation (but not phosphorylation of other MPKs) could be detected in un-elicited seeds (Figure 6b, Supplementary Figure 16). Following quantification, we found that the degree of phosphorylation of MPK6 was reduced by approximately 50% in *gso1-1 gso2-1* double mutant seeds compared to wild-type, suggesting that a significant proportion of MPK6 phosphorylation in seeds depends on the activity of GSO1 and GSO2 (Figure 6c, Supplementary Figure 16). Intriguingly, in seeds, a band corresponding to a second phosphorylated MPK was detected exclusively in *mpk6-2* mutants (Figure 6b), suggesting that the relatively weak *mpk6* seedling cuticle phenotype could be due to compensation by an as yet unidentified MPK [53].

### MPK6 activity is required in the embryo, but not the endosperm, to maintain cuticle integrity

The strong expression of GSO1 and GSO2 in the embryonic epidermis, suggests that the activity of GSO1 and GSO2 in cuticle formation is required in the embryo. No promoters confirmed as specifically being expressed only in the embryo or embryo epidermis, have been published. To further confirm the spatial requirement for GSO1/GSO2-dependent signalling in the seed, we therefore complemented the *mpk6-2* mutant either with the *MPK6* cDNA expressed under the ubiquitously expressed *RPS5A* promoter, or under the endosperm specific *RGP3* promoter [54,55]. We were unable to complement either the misshapen seed/seed bursting phenotypes or the toluidine blue permeability phenotypes of *mpk6-2* mutants by expressing *MPK6* in the endosperm, but obtained full complementation of all phenotypes in plants expressing *MPK6* under the *RPS5A* promoter (Figure 7, Supplementary Figures 17). Together with the results of our reciprocal crosses, these findings indicate that the seedling permeability phenotype of *mpk6-2* mutants is most likely due to signalling defects in the embryo. Seed size and seed bursting defects could be caused by lack of MPK6 in the testa, as suggested by reciprocal crosses, although this remains to be investigated in more detail. In order to further confirm the function of MPK6 downstream of GSO1/GSO2 signalling we attempted to express a constitutively active form of MPK6 under the *RPS5A* promoter in wild type and double mutant plants, but were unable to generate any transformants, potentially due to the critical roles played by MPK6 during early embryogenesis.

**Figure 7.**
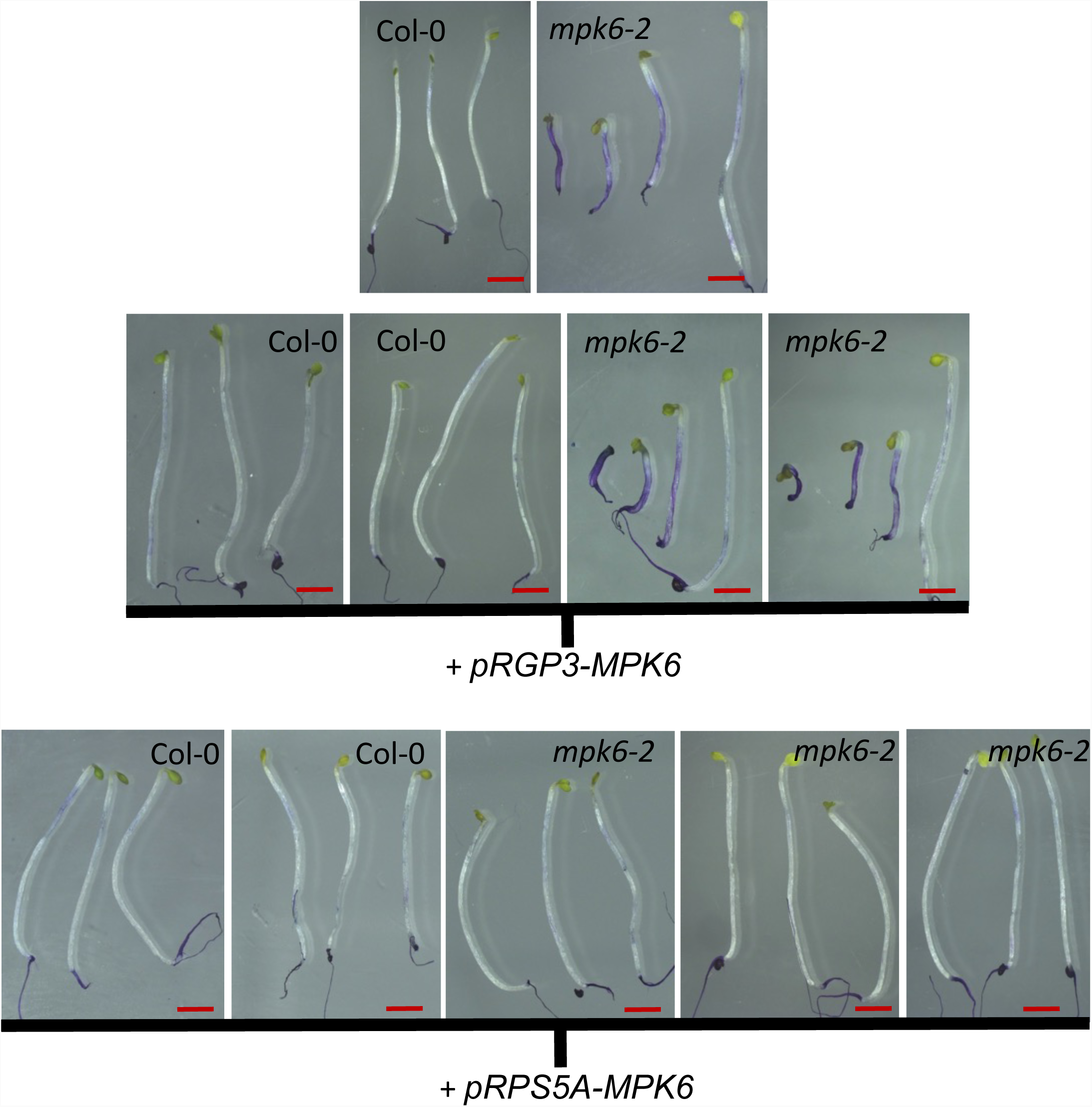
MPK6 activity is required in the embryo and testa, but not the endosperm, for normal seedling development. (a) Representative phenotypes of toluidine blue-stained seedlings from wild-type (Col-0), *mpk6-2*, and these backgrounds transformed with *pRGP3-MPK6* or *pRPS5A-MPK6.* Lines correspond to those described in Supplementary Figure 17. Scale bar = 2mm

## DISCUSSION

It this study, consistent with the similarity between GSO1/2 and PEPR1/2 proteins, we found that stress-associated kinase MPK6, which has been shown to act downstream of PEPR signalling [56], shows constitutive phosphorylation in developing seeds, and that this phosphorylation is partially dependent upon GSO1 and GSO2. In addition, we showed that GSO1/GSO2, are required for the expression of a set of stress-related genes during early seed development. Our results suggest that GSO1/GSO2 dependent stress response-related signalling pathways are active in developing seeds. Because of the conserved transcriptional targets expressed downstream of GSO1/GSO2 dependent signalling, and in defence responses, this scenario is distinct from previously reported situations in which single pathway components, such as the co-receptor BAK1, play distinct roles in developmental and defence-related signalling cascades through interaction with multiple receptors [57,58]. However, the role of the transcriptional targets of GSO1/GSO2 signalling in seeds remains to be elucidated.

Our work also shows that GSO1/GSO2, ALE1 and MPK6 act in a genetic pathway involved in ensuring embryonic cuticle integrity. We show for the first time that embryonic cuticle biogenesis involves the coalescence of discontinuous patches of cutin-like material that appear on the embryo surface at the globular stage, and that pathway mutants are either incapable of completing, or retarded in the completion of “gap closure” during this process. Interestingly, GSO1 (also known as SCHENGEN3 [6]) was recently shown to be involved in ensuring the continuity of another apoplastic diffusion barrier, the Casparian strip, which prevents the apoplastic movement of solutes from the cortex to the stele of the root [6]. GSO1 may therefore form part of a general mechanism employed by plants for monitoring the “integrity” of apoplastic barriers formed during plant development.

The role of GSO1 and GSO2 in the closure of gaps in the nascent cuticle implies spatial regulation of signalling outputs at the subcellular level. Cytoplasmic signalling components which, like MPK6 might not be uncovered by transcriptome analysis but rather are modified post-translationally, are therefore likely to be of critical importance in GSO1 GSO2 signalling in the embryonic epidermis. Indeed, although MPK6-mediated signalling has most often been implicated in the control of transcription, particularly via the modulation of the activity of WRKY transcription factors, evidence for potential roles in cytoplasmic responses, for example during funicular guidance of pollen tubes [46] and control of cell division planes [50], exist in the literature.

Cytoplasmic responses downstream of receptor-like kinases include the local production of apoplastic Reactive Oxygen Species (ROS) and/or calcium influxes, and indeed localized ROS production has been implicated in Casparian strip formation [6,59-61]. However although a plausible model has proposed that ROS release could mediate Casparian strip polymerisation though polymerisation of monolignols [59], it is less obvious how ROS could directly affect the biosynthesis of an aliphatic cutin-based barrier, although a possible role for ROS in linking the cuticle to the cell wall has been evoked [62]. ROS production has been shown to directly modulate the activation of MAPK signalling, providing a mechanism permitting the reinforcement of localised signalling events [63,64]. Another, potentially linked, possibility is that GSO1/GSO2 activity in the embryo could spatially direct the secretion of either cuticle components or enzymes and cell wall components necessary for their integration into the cutin polymer, in a system analogous to the rapid and highly localized deposition of callose observed upon hyphal penetration into epidermal cells (reviewed in [65,66]). Interestingly MPK6 has also been shown to be involved in phragmoplast formation during root cell division and therefore could be involved in the localised production/secretion of apoplastic compounds [50]. However, observing these processes *in situ*, within the living seeds, would require developments in microscopy which are not yet available.

Our work highlights several questions which merit further discussion. A first important question is whether the GSO1/GSO2 signalling pathway could play a role in protecting seeds, or more generally plants, against pathogen attack. Cuticle integrity in adult plants has been shown to be required for resistance to *Pseudomonas* pathovars [67,68]. The action of the ALE1 GSO1/GSO2 signalling pathway in ensuring embryonic cuticle integrity is therefore likely to have a significant influence on embryo and seedling susceptibility to bacterial pathogens. However, we have also shown that GSO1/GSO2, ALE1 and MPK6 are necessary for the expression of known defence marker genes in seeds. Cuticle permeability phenotypes have neither been reported in the literature for mutants affected in the defence markers identified in our transcriptome studies, nor found in our own studies (unpublished results). This raises the question of whether the ALE1, GSO1/GSO2, MPK6 signalling pathway, in addition to mediating localised apoplastic modifications, could act at a more global level either to protect developing seeds from the ingress of bacterial pathogens (thus affecting vertical pathogen transmission), or to “prime” embryos against pathogen attack upon germination. Exploring this possibility would necessitate functionally separating susceptibility caused by cuticle defects from lack of immune priming, and will be technically very challenging, but could ultimately inform strategies aiming to reduce vertical transmission of plant pathogens.

The second question concerns how signalling via GSO1 and GSO2 is triggered in the seed in the absence of pathogens. In this study we consolidate data supporting the function of ALE1 in the same pathway as GSO1 and GSO2. We previously proposed that the function of ALE1, GSO1 and GSO2 in ensuring the apoplastic separation of the embryo and endosperm became necessary in angiosperms due to developmental constraints imposed by the sexualisation of the female gametophyte, which led to the simultaneous development of the embryo and surrounding nutritive tissues post-fertilization, rather than their sequential development [11,12]. *ALE1* expression is endosperm specific and, as previously suggested [69], the recruitment of *ALE1* to a function in reinforcing the embryonic cuticle may have occurred during the emergence of the angiosperm lineage. Subtilases have been shown to be involved in defence responses and immune priming in plants [70,71]. It is thus possible that ALE1 acts to produce an as yet unidentified ligand for the GSO1 and GSO2 receptors. In such a scenario the function of ALE1 in the seed could be analogous to the “immune priming” function previously reported for the subtilase SBT3.3 [71].

Such a scenario naturally raises a third, and important question, around the identity of the ligand of GSO1 and GSO2. Two sulfated peptides, CIF1 and CIF2, which can act as ligands for GSO1 during Casparian strip formation, have recently been identified [72,73]. Testing the role of these molecules in developing seeds will be an obvious priority. However Nakayama and colleagues specifically reported that no cuticle defects (as gauged by cotyledon fusion phenotypes) were observed in *cif1 cif2* double mutants and the possibility that other signalling molecules could be involved in ensuring embryonic cuticle integrity therefore cannot be excluded.

In summary, we propose that endosperm-localised factors (like ALE1) may have been recruited to hijack a defence-signalling pathway involving the ancestor(s) of GSO1 and GSO2, and downstream signalling components including MPK6, and trigger an “auto-immune” type response in the embryo to ensure cuticle integrity. The future identification of further pathway components, and in particular the substrates of ALE1 and ligands of GSO1 and GSO2, will help to confirm this hypothesis.

## Materials and Methods

### Plant material

The *pGSO1*:GSO1-mVENUS line was kindly donated by Professor Niko Geldner (Unil-Sorge, University of Lausanne). The *mpk6-2 (*SALK_073907) mutant and the *mpk3-1 (*SALK-151594) were kindly provided by Dr Roberta Galletti.

### Growth conditions and plant treatments

Unless otherwise specified, plants were grown for 10 d in sterile conditions on Murashige and Skoog (MS) agar plates with 0.5% sucrose, 1 month under short-day conditions (19°C, 8h light / 17°C, 16h dark) and then transferred to standard long-day conditions (21°C, 16h light/8h dark) for one more month. To stage material, newly opened flowers were marked each day for two weeks. For bacterial growth assays, plants were grown under controlled conditions in a growth chamber at 21°C, with a 9-hours light period and a light intensity of 190 μmol.m^−2^.s^−1^. For MPK6 activation analysis, seedlings were grown for 10 d in MS liquid medium supplemented with 0.5% sucrose in 100μm cell strainers submerged in 6-well plates (5ml of medium per well). Cell strainers were transferred to new plates containing MS sucrose 0.5% supplemented with 100 nM or 1 nM flg22 or water and incubated for 15 and 60 minutes at room temperature without skaking. Seedlings were then rapidly harvested in liquid nitrogen and stored at −80 °C until protein extraction. For cutin analysis of seedlings, seeds were sterilized, plated on MS medium supplemented with 0.7 % agar, 0.7 % sucrose and 2.5 mM MES-KOH, pH 5.7, and stratified in the dark for 3 days at 4°C. Plates were then transferred to a controlled environment growth chamber at 22°C and with continuous light, and seedlings were grown for 5 days before harvesting the cotyledons. For toluidine blue staining and Nile-Red staining, sterilized seeds were spread uniformly on 15 cm MS plates with 0.5% sucrose and 0.4% Phytagel (Sigma) (pH 5.8) and stratified for 2 days in the dark at 4°C. After stratification seeds were transferred to a growth chamber and incubated for 6h under continuous light followed by 4 days in the dark.

### *In situ* hybridization

DNA templates for the probes used in *in situ* hybridizations were amplified using the primers listed in Supplementary Table 2. Digoxigenin-labelled RNA probes were produced and hybridized to tissue sections following standard procedures. In brief, siliques were opened, fixed overnight in ice-cold PBS containing 4% paraformaldehyde, dehydrated through an ethanol series, embedded in Paraplast Plus (Mc Cormick Scientific) and sectioned (8 μm). Immobilized sections were dewaxed and hydrated, treated with 2x saline sodium citrate (20 min), digested for 15 min at 37°C with proteinase K (20 mg/ml) in 50mM Tris-HCl, pH 7.5, 5mM EDTA), treated for 2 min with 0.2% glycine in PBS, rinsed, postfixed with 4% paraformaldehyde in PBS (10 min, 4°C), rinsed, treated with 0.25% w/v acetic anhydride in 100mM triethanolamine (pH 8.0 with HCl) for 10 min, rinsed and dehydrated. Sections were then hybridized under coverslips overnight at 50°C with RNA probes (produced using DIG RNA labelling kit (Roche)) diluted in DIG easy Hyb solution (Roche) following the manufacturer’s instructions. Following hybridization, the slides were extensively washed in 0.1x saline sodium citrate and 0.5% SDS at 50 °C (3 h), blocked for 1 hour in 1% blocking solution (Roche) in TBS and for 30 minutes in BSA solution (1% BSA, 0.3% Triton-X-100, 100mM Tris-HCl, 100mM NaCl, 50mM MgCl_2_), and then incubated in a 1/3000 dilution of in alkaline phosphatase-conjugated antidigoxigenin antibody (Roche) in BSA solution for 2 h at RT. Sections were extensively washed in BSA solution, rinsed and treated overnight in the dark with a buffered NBT/BCIP solution. Samples were rinsed in water before air drying and mounting in Entellan (Sigma).

### Microscopy

Embryos were imaged by gently bursting seeds between slide and cover-slip in water and imaging using a dipping lens with a long working distance. Confocal imaging was carried out on a Zeiss LSM700 with a W N-Acroplan 40x/0.75 M27 objective. mVENUS was excited using a 488nm diode laser and fluorescence was collected using a 490-555 nm PMT. Light microscopy imaging was carried out using a Zeiss axioimager 2. Images were acquired using bright field illumination.

### Histochemical staining with Nile-Red

5-day-old etiolated seedlings were stained with Nile-Red (Sigma-Aldrich, stock solution at 1mg/mL in DMSO) at 2μg/mL in 50mM PIPES (Sigma-Aldrich) pH 7.0. After 20min of incubation in dark, seedlings were washing 3-times in water and placed between slide and lamella. Confocal imaging was performed using a Zeiss LSM700 with 488nm excitation and >530nm emission filters. Images were then processed in the Zeiss LSM Image Browser Program.

### Cutin analysis

Cuticle composition and content was analyzed as previously described [74,75].

### TEM analysis

For transmission electron microscopy analysis, seeds were removed from siliques by removal of the replum tissue with attached seeds. Seeds were high-pressure frozen with a Leica EM-PACT-1 system. Three seeds were inserted into a flat copper carrier, fast-frozen, and cryosubstituted into the Leica AFS1 device. The different freeze-substitution steps were as follows: 54 h at −90°C in acetone solution containing 0.2% glutaraldehyde, 1% osmium tetroxide, and 0.1% uranyl acetate. The temperature was then raised with a step of 2°C/h before remaining for 8 hours at −60°C. The temperature was raised again to −30°C for 8h00 before being increased to 4°C. Samples were washed three times for 10 min in 100% acetone before embedding in Spurr’s resin, which was performed progressively (8 h in 25% Spurr’s resin in acetone, 24 h in 50% Spurr’s resin in acetone, 24 h in 75% Spurr’s resin in acetone, and two times for 12 h in 100% Spurr’s resin). Polymerization was performed at 70°C for 18 h.

Samples were sectioned (65 nm sections) and imaged at 120 kV using an FEI TEM tecnai Spirit with 4 k × 4 k eagle ccd.

### Generation of micro-array data

Microarray analysis was carried out at a Transcriptomic Platform, POPS, at the Institute of Plant Sciences Paris-Saclay (IPS2, Orsay, France), using a CATMAv7 array based on AGILENT technology [76]. The CATMAv7 array for the Arabidopsis thaliana genome was made using gene annotations included in FLAGdb++, an integrative database of plant genomes (http://urgv.evry.inra.fr/FLAGdb, [77]). The single high density CATMAv7 microarray slide contains four chambers, each containing 149 916 primers. Each 60 bp primer is present in triplicate in each chamber for robust analysis, and as both strands. The array contains 35 754 probes (in triplicate) corresponding to genes annotated in TAIRv8 (among which 476 probes correspond to mitochondrial and chloroplast genes), 1289 probes corresponding to EUGENE software predictions, 658 probes to miRNA/MIRs and 240 control probes.

3 independent biological replicates were produced. For each biological repetition and each point, RNA samples were obtained by pooling RNAs from staged siliques containing embryos at the pre-globular to globular, or the young to late heart stage. Total RNA was extracted using the Spectrum™ Plant Total RNA Kit (Sigma-Aldrich) according to the suppliers’ instructions. For each comparison, one technical replicate with fluorochrome reversal was performed for each biological replicate (i.e. four hybridizations per comparison). The labelling of cRNAs with Cy3-dUTP or Cy5-dUTP was performed as described in the Two-Color Microarray-Based Gene Expression Analysis Low Input Quick Amp Labeling manual (© Agilent Technologies, Inc.). The hybridization and washing steps were performed according to the Agilent Microarray Hybridization Chamber User Guide instructions ((© Agilent Technologies, Inc.). Two micron scanning was performed with InnoScan900 scanner (Innopsys^R^, Carbonne, FRANCE) and raw data were extracted using Mapix^R^ software (Innopsys^R^, Carbonne, FRANCE).

### Statistical Analysis of Microarray Data

Experiments were designed with the statistics group of the Unité de Recherche en Génomique Végétale. For each array, the raw data comprised the logarithm of median feature pixel intensity at wavelengths 635 nm (red) and 532 nm (green). For each array, a global intensity-dependent normalization using the loess procedure [78] was performed to correct the dye bias. The differential analysis was based on log-ratio averaging over the duplicate probes and over the technical replicates. Hence the number of available data points for each gene equals the number of biological replicates and is used to calculate the moderated t-test [79]. Analysis was carried out using the R software (http://www.R-project.org). Under the null hypothesis, no evidence that the specific variances vary between probes was highlighted by Limma and consequently the moderated t-statistic was assumed to follow a standard normal distribution. To control the false discovery rate, adjusted p-values found using the optimized FDR approach [80] were calculated. We considered as being differentially expressed, the probes with an adjusted p-value ≤ 0.05. The function SqueezeVar of the library Limma was used to smooth the specific variances by computing empirical Bayes posterior means. The library kerfdr was used to calculate the adjusted p-values.

### Data Deposition

Microarray data from this article were deposited at Gene Expression Omnibus (http://www.ncbi.nlm.nih.gov/geo/), accession no. GSE68048) and at CATdb (http://urgv.evry.inra.fr/CATdb/; Project: AU14-04_INASEED) according to the “Minimum Information About a Microarray Experiment” standards.

### Quantitative gene expression analysis in seeds

Intact siliques were frozen in liquid nitrogen and total RNA was extracted using the Spectrum Plant Total RNA Kit (Sigma). Total RNAs were digested with Turbo DNA-free DNase I (Ambion) according to the manufacturer’s instructions. RNA was reverse transcribed using the SuperScript VILO cDNA Synthesis Kit (Invitrogen) according to the manufacturer’s protocol. PCR reactions were performed in an optical 384-well plate in the QuantStudio 6 Flex System (Applied Biosystems), using FastStart Universal SYBR Green Master (Rox) (Roche), in a final volume of 10 μl, according to the manufacturer’s instructions. The following standard thermal profile was used for all PCR reactions: 95 °C for 10 min, 40 cycles of 95 °C for 10 s, and 60 °C for 30 s. Data were analysed using the QuantStudio 6 Flex Real-Time PCR System Software (Applied Biosystems). As a reference, primers for the EIF4A cDNA were used. PCR efficiency (E) was estimated from the data obtained from standard curve amplification using the equation E=10^−1/slope^. Expression levels are presented as 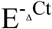, where ΔCt=Ct_GOI_-Ct_EIF4A_. Primers are listed in Supplementary Table 2.

### Toluidine blue staining

The lids of plates containing etiolated seedlings were removed and plates were immediately flooded with staining solution [0.05% (w/v) Toluidine Blue + 0.4% (v/v) Tween-20] for 2 minutes. The staining solution was poured off and plates were immediately rinsed gently by flooding under a running tap until the water stream was no longer visibly blue (1-2 minutes). Seedlings were photographed under a Leica MZ12 stereomicroscope.

### Protein extraction and MPK6 activation analysis

Seedlings or seeds were quickly frozen in liquid nitrogen and proteins were extracted in buffer containing 50 mM Tris pH 7.5, 200 mM NaCl, 1 mM EDTA pH 8, 10% glycerol, 0.1% tween 20, 1 mM phenylmethylsulfonyl fluoride, 1 mM dithiothreitol, 1x protease inhibitor cocktail P9599 (Sigma-Aldrich), and 1x MS-Safe protease and phosphatase inhibitor cocktail (Sigma-Aldrich). Equal amounts of proteins (20 μg for seedlings and 10 μg for seeds) were resolved on 10% polyacrylamide gels and transferred onto a nylon membrane (Schleicher & Schuell). For seedlings primary antibodies against phospho p44/42 MAP kinase (1:2000 dilution) (Cell Signaling Technologies) and then against MPK6 (1:10000 dilution) (Sigma-Aldrich) were used with horseradish peroxidase-conjugated anti-rabbit as secondary antibody. Signal detection was performed using the SuperSignal™ West Femto Maximum Sensitivity Substrate kit (Pierce). For seeds primary antibodies against phospho p44/42 MAP kinase and then against MPK6 were used with IRDye® 800CW Donkey anti-Rabbit IgG (H + LI-COR, 1:10000 dilution), and the bound complex was detected using the Odyssey Infrared Imaging System (Li-Cor; Lincoln, NE). The images were analysed and quantified with ImageJ. Background was subtracted for each band. To test the linearity of the detection, 5-15 μg protein from heart stage developing seeds were treated as previously. To detect the antibody against phospho p44/42 MAP kinase an anti-Rabbit IgG, HRP conjugate (Amersham, 1:30000) was used. Anti-alpha-tubulin (Sigma, 1:2000) was used with an anti-mouse IgG, HRP conjugate (GE HealthCare, 1:10000). Signal detection was performed using Clarity Max™ Western ECL Substrate (Biorad) with a ChemiDoc Touch (Biorad) instrument. The images were quantified with ImageJ. Background was subtracted for each band.

## Acknowledgements

We would like to thank Professor N. Geldner for providing seeds, and A. Lacroix, J. Berger, P, Bolland, H. Leyral and I. Desbouchages for assistance with plant growth and logistics. AC was funded by a grant from the French National Research Agency (ANR-13-BSV2-0002, INASEED). SM was supported by a doctoral grant from the Rhône-Alpes region. RG was supported by a European Research Council Starting Grant (Phymorph #307387). NMD is funded by a PhD fellowship from the Ministère de l’Enseignement Supérieur et de la Recherche. Microscopy and lipid analyses were respectively performed at the Bordeaux Imaging Center (which is a member of the national infrastructure France BioImaging), and the Metabolome Facility of the Functional Genomic Center of Bordeaux (which is supported by the grant MetaboHUB-ANR-11-INBS-0010).

## Author contributions

Experiments were carried out by AC, LB, JJ, LT, NMD, SP, SM, ACM and FD. Results were analyzed by all authors. GI, FD, JJ, TW and RG designed experiments and supervised the work. GI wrote the paper with contributions from all authors.

**The authors declare that they have no competing financial interests.**

## Supporting Information Legends

**Supplementary Figure 1 (Related to Figure 1) : Genes involved in cuticle biosynthesis are expressed early during embryo development and are co-expressed with *GSO1* and *GSO2*.** Expression data for *LACS2 (***a)**, *FDH (***b)**, *BDG (***c)**, *LCR (***d)**, *LTPG1 (***e)**, *ABCG11 (***f)**, *GSO1 (***g)** and *GSO2 (***h)** downloaded from the Seed Gene Network resource (http://seedgenenetwork.net/).

**Supplementary Figure 2 (Related to Figure 1) : Genes involved in cuticle biosynthesis are co-expressed with *GSO1* and *GSO2* in the embryonic epidermis during embryogenesis, but their expression is not dependent upon GSO1 and GSO2.** Analysis of the expression of genes involved in cuticle biosynthesis in wild-type (Col-0) and *gso1-1 gso2-1* seeds containing late globular/triangle, heart and early torpedo stage embryos (left to right).

**Supplementary Figure 3 (Related to Figure 1) : Hybridization of tissue sections to an antisense *GFP* probe as negative control in Col-0 (a-c) and *gso1 gso2 (*d-f) seeds containing late globular (a,d), heart (b,e) and early torpedo (c,f) stage embryos.** Expression of *LTPG1* in wild type seed containing torpedo stage embryo is shown in (g) for direct comparison.

**Supplementary Figure 4 (Related to Figure 4) : Analysis of the permeability of extruded Col-0 embryos at different developmental stages to toluidine blue treatment.** Three stained and one unstained embryo (below) are shown for each developmental stage. Scale bar = 100mm

**Supplementary Figure 5 (Related to Figure 4) : Cuticle discontinuities in early wild-type (Col-0) embryos and later *gso1 gso2* embryos. a,b)** Cuticle the mid-late globular stage (during gap closure) in Col-0 embryos showing disconuinuous cuticle. *gso1-1 gso2-1* mutant embryos maintain a diffuse and discontinuous cuticle at later stages. Analysis of embryonic cuticle deposition in *gso1-1 gso2-1* at the *mid heart (***c-d)** and walking stick (**e-f)** stages of embryogenesis, when wild-type cuticle is continuous. White arrows show external face of embryonic cuticle. Scale bar = 500nm.

**Supplementary Figure 6 (Related to Figure 5) : GO term analysis of genes showing increased expression in siliques of both *gso1-1 gso2-1* and *ale1-4* mutant backgrounds at the globular (a) and heart (b) stages of embryo development.** The degree of overlap between these datasets is illustrated in (**c)**.

**Supplementary Figure 7 (Related to Figure 5) : GO term analysis of genes showing reduced expression in siliques of both *gso1-1 gso2-1* and *ale1-4* mutant backgrounds at the globular (a) and heart (b) stages of embryo development.**

**Supplementary Figure 8 (Related to Figure 5) : Validation by qRT-PCR of microarray data.** Experiments were carried out in 4 biological replicates. Values are expressed relative to the *EIF4A* gene. Significance values indicated were calculated using a Student’s t-test. *** denotes p<0.01, ** denotes p<0.05 and * denotes p<0.1. Bars indicate standard errors.

**Supplementary Figure 9 (Related to Figure 5) : Analysis of the expression of *SWI3A* in wild-type (a-c), *gso1-1 gso2-1 (*d-f) mutant embryos in seeds containing late globular/triangle (a,d), heart (b,e) and early torpedo (c,f) stage embryos.** Expression data for *SWI3A* downloaded from the Seed Gene Network resource (http://seedgenenetwork.net/) is shown in (**g).**

**Supplementary Figure 10 (Related to Figure 6) : Non additivity of seed twisting (a-b) and seedling cuticle permeability (c) phenotypes between *mpk6-2* and *ale1-4* mutants and between *mpk6-2* and *gso1-1 gso2-1* double mutants.** Populations of seeds (**a)** from single double and triple mutants were photographed, and seed phenotypes were quantified (**b)**(Col-0 n= 196, *ale1-4* n=200, *mpk6-2* n=210, *gso1-1 gso2-1* n=111, *mpk6-2 ale1-4 (*3 individuals) n = 211, 212 and 238, *mpk6-2 gso1-1 gso2-1* n=86). Etiolated seedlings were treated with toluidine blue and seedling and toluidine blue phenotypes were quantified (**c)**. Results are representative of three independent experiments. Col-0 n=389, *ale1-4* n=387, *mpk6-2* n=383, *mpk6-2 ale1-4* n=398. (Quantifications were not possible for *mpk6-2 gso1-1 gso2-1* triple mutants due to low seed set).

**Supplementary Figure 11 (Related to Figure 6) : Cuticle phenotypes using Nile-Red staining of etiolated cotyledons.** Genotypes are indicated on left panels with zones magnified on the right highlighted by white boxes. Arrows indicate position of the cuticle and arrowheads indicate gaps in the cuticle. a) Biological replicate 1.

**Cuticle phenotypes using Nile-Red staining of etiolated cotyledons.** b) Biological replicate 2

**Supplementary Figure 13 (Related to Figure 6) : qRT-PCR analysis of the expression of candidate target genes in *mpk6-2* mutant siliques.** Experiments were carried out in biological triplicate. Values are expressed relative to *EIF4* gene expression. Significance values indicated were calculated using a Student’s t-test. *** denotes p<0.01, **denotes p<0.05 and * denotes p<0.1. Bars indicate standard errors.

**Supplementary Figure 14 (Related to Figure 6) : GSO1 and GSO2 are not necessary for MPK6 phosphorylation in response to PAMP-elicitation in seedlings.** Western blot analysis of phosphorylated MPK proteins (upper panel) and then total MPK6 protein (middle panel). Loading control (Ponceau S-stained Rubisco) is shown in lower panel. The same blot is shown in the upper middle and lower panel. * Indicates a non specific band detected by the anti-MPK6 antibody. Seedlings were treated with water or with 100nM flg22 for either 15 or 60 minutes before protein extraction. The mutants *mpk3-1* and *mpk6-2* were included to confirm band identities.

**Supplementary Figure 15 (Related to Figure 6) : *gso1 gso2* mutant seedlings are not significantly affected in the MPK6 phosphorylation response to flg22.** Western blot analysis of phosphorylated MPK proteins (upper panels) and then total MPK6 protein (middle panels). Loading control (Ponceau S-stained Rubisco) is shown in lower panel. The same blot is shown in the upper middle and lower panel. * Indicates a non specific band detected by the anti-MPK6 antibody. Seedlings were treated with water or with 1nM flg22 for either 15 or 60 minutes before protein extraction.

**Supplementary Figure 16 (Related to Figure 6) : Developing *gso1 gso2* mutant seeds show reduced levels of MPK6 phosphorylation compared to wild-type seeds.** a) Western blot analysis of phosphorylated MPK6 protein in developing seeds exposed at four consecutive one minute intervals (to confirm signal linearity). Loading control (α-tubulinA) is shown in lower panel. B) Degree of phosphorylation of MPK6 in Col-0 and *gso1-1 gso2-1* mutant seeds. Error bars represent SD of 3 biological replicates.

**Supplementary Figure 17 (Related to Figure 7) : MPK6 is required in the embryo and testa, but not the endosperm (a)** Representative phenotypes of seeds from wild-type (Col-0), *mpk6-2*, and these backgrounds transformed with *pRGP3-MPK6 or pRPS5A-MPK6. (***b)** Quantification of seed phenotypes in the above material. Seeds from at least two independent transgenic lines have been quantified.

